# Deep learning for automated analysis of fish abundance: the benefits of training across multiple habitats

**DOI:** 10.1101/2020.05.19.105056

**Authors:** Ellen Ditria, Michael Sievers, Sebastian Lopez-Marcano, Eric L. Jinks, Rod M. Connolly

## Abstract

Environmental monitoring guides conservation, and is thus particularly important for coastal aquatic habitats, which are heavily impacted by human activities. Underwater cameras and unmanned devices monitor aquatic wildlife, but manual processing of footage is a significant bottleneck to rapid data processing and dissemination of results. Deep learning has emerged as a solution, but its ability to accurately detect animals across habitat types and locations is largely untested for coastal environments. Here, we produce three deep learning models using an object detection framework to detect an ecologically important fish, luderick (*Girella tricuspidata*). Two were trained on footage from single habitats (seagrass or reef), and one on footage from both habitats. All models were subjected to tests from both habitat types. Models performed well on test data from the same habitat type (object detection measure: mAP50: 91.7 and 86.9% performance for seagrass and reef, respectively), but poorly on test sets from a different habitat type (73.3 and 58.4%, respectively). The model trained on a combination of both habitats produced the highest object detection results for both tests (92.4 and 87.8%, respectively). Performance in terms of the ability for models to correctly estimate the ecological metric, MaxN, showed similar patterns. The findings demonstrate that deep learning models extract ecologically useful information from video footage accurately and consistently, and can perform across habitat types when trained on footage from the variety of habitat types.

## Introduction

People have been monitoring and counting wildlife for millennia, collecting invaluable data for a number of uses such as informing conservation, tracking population trends and estimating abundance or biomass for fisheries stock assessments (Goldsmith 2012). As the world changes and many terrestrial and marine ecosystems experience severe and sustained declines in extent and condition (Maxwell et al. 2016), monitoring wildlife has never been more important. The speed and scale at which the natural world is changing also mean that monitoring and analysing data quickly enough to be able to respond has become a global challenge. Aquatic coastal habitats are among the most severely affected by anthropogenic activities (Davidson 2014; Tulloch et al. 2020), despite being renowned for their roles in fisheries productivity, coastal protection, carbon sequestration and biodiversity (Sievers et al. 2019; Silliman et al. 2019). Although the challenge of developing rapid, effective monitoring is important in all environments, monitoring wildlife in coastal aquatic habitats is particularly important.

The advent of cheap, high-resolution cameras has enabled large amounts of underwater data to be collected without many of the logistical issues encountered using traditional, manual methods of data collection. For example, cameras can be deployed in situ for periods of hours to months to collect data without the need for human interaction (Podder et al. 2019). Additionally, the presence of humans and their equipment often causes animals to display avoidance behaviour and make data collection unreliable (Frid and Dill 2002). The ease with which data can now be collected, however, has only exacerbated the challenge of being able to analyse the data quickly, with manual analysis of photo and video footage laborious (Weinstein 2018). Scientists consequently need tools to analyse enormous amount of monitoring data and quickly extract useful ecological information for management and conservation purposes.

Deep learning technologies have emerged as elegant solutions for automating the analysis of video and image-based datasets. Deep learning is a derivative of machine learning, which is broadly categorised as algorithms that can generate a prediction based on pattern detection in data (Christin et al. 2019). This technology outperforms traditional machine learning algorithms which are limited in their ability to process raw images as they often require manual feature extraction prior to data analysis (LeCun et al. 2015). Further, deep learning algorithms have greater accuracy than traditional machine learning algorithms when presented with underwater imagery (Villon et al. 2016). Scientists have recently utilised deep learning technology to detect and identify fish to the species level (Villon et al. 2018; dos Santos and Goncalves 2019; Salman et al. 2019b), and to count individual fish of a target species in video frames to provide estimates of MaxN (Ditria et al. 2019). Importantly, these methods can perform at higher accuracies and speeds than humans (Ditria et al. 2019). Although this allows scientists to greatly increase the efficiency, repeatability, and accuracy of image-based data analysis (Weinstein 2018), training these algorithms takes a considerable amount of time and imposes high initial costs (Christin et al. 2019). Flexibility and robustness in identifying species across large spatial and temporal scales are therefore key for deep learning to be a worthwhile method to replace manual analysis

The use of deep learning to monitor and count wildlife such as fish in coastal environments also presents a unique set of challenges. For example, there are many factors that may affect the model’s ability to detect fish, such as water turbidity, lighting variation, occlusion due to schooling fish, and changes in fish orientation (Mandal et al. 2018). Further, many fish utilise a variety of different habitats, whether it is daily migrations among habitats to feed and shelter, or ontogenetic habitat shifts (Lecchini and Galzin 2005; Igulu et al. 2014). Among these different habitats, there may be large variations in local fish assemblages. Since deep learning models are trained with footage, these inter-habitat differences might mean that a singularly trained deep learning model may not be reliable across habitats. For instance, structural complexity of the habitat may influence the performance of the model, as background confusion and foreground camouflage might compromise the model’s accuracy (Salman et al. 2019a). Accounting for different video and image backgrounds is a challenging problem in computer science, especially in real-world footage that displays significant lighting changes and complex and dynamic backgrounds, such as those from underwater environments (Dou et al. 2019).

To maximise the effectiveness of monitoring species, as well as the reliability of analysis, deep learning models must be robust across multiple habitats that target species utilises. Do date, however, it is not known how well deep learning algorithms perform across multiple habitat sites, or if a model trained on a singular habitat will perform as well when tested on another (e.g. training with seagrass footage and tested on reef footage). Here, we test the potential for deep learning algorithms to work effectively across habitats using an ecologically and economically important fish species, luderick (*Girella tricuspidata*). Luderick are found in multiple habitats including seagrass meadows and rocky reefs along the temperate waters of east coast Australia and northern New Zealand (Abrantes et al. 2015). They are a recreational and commercial fisheries species, and are important herbivores that control algal growth on reefs and prevent smothering of seagrass by epiphytic algae (Ferguson et al. 2015). By assessing the capacity of deep learning algorithms to transcend habitats, we provide evidence of the applicability of this technology to assist monitoring and conservation efforts.

## Methods

### Datasets

The training data set was collected using submerged action cameras (Haldex Sports Action Cam HD 1080p, GoPro 8 Black 1080p) deployed in the two most dissimilar habitats frequented by luderick, seagrass meadows and rocky reefs, between February 2019 and March 2020, in the Tweed River estuary, northern New South Wales, Australia (−28.169438, 153.547594). Cameras were positioned to collect footage at a range of angles and backgrounds to ensure variety in the training data. Footage was also collected over several days to increase variability in other environmental factors such as lighting and water turbidity. Videos were trimmed to remove footage with frames empty of fish and split into 5 frames per second. Polygonal segmentation masks were manually drawn around the region of interest (ROI), here individual luderick, and were only annotated if they could be positively identified by an expert fish taxonomist at any time within the video. The algorithm then extracts features automatically and begins to recognise patterns which “train” the computer to associate these with the ROI (LeCun et al. 2015). Three separate datasets, each consisting of ∼ 4,700 annotated luderick, were used for training containing: seagrass footage only, reef footage only, and a combination of both habitats using half the annotations of the single habitat training datasets (Figure 1). The two test datasets in the different habitat types (seagrass and reef) were comprised of video footage that did not appear in the training data (Figure 1). The two test sets each comprised of 48 videos with approximately 1,500 luderick annotations which were pre-annotated and used as the ground truth to quantify the model’s ability to accurately detect and count fish (Figure 1).

**Fig. 1.**
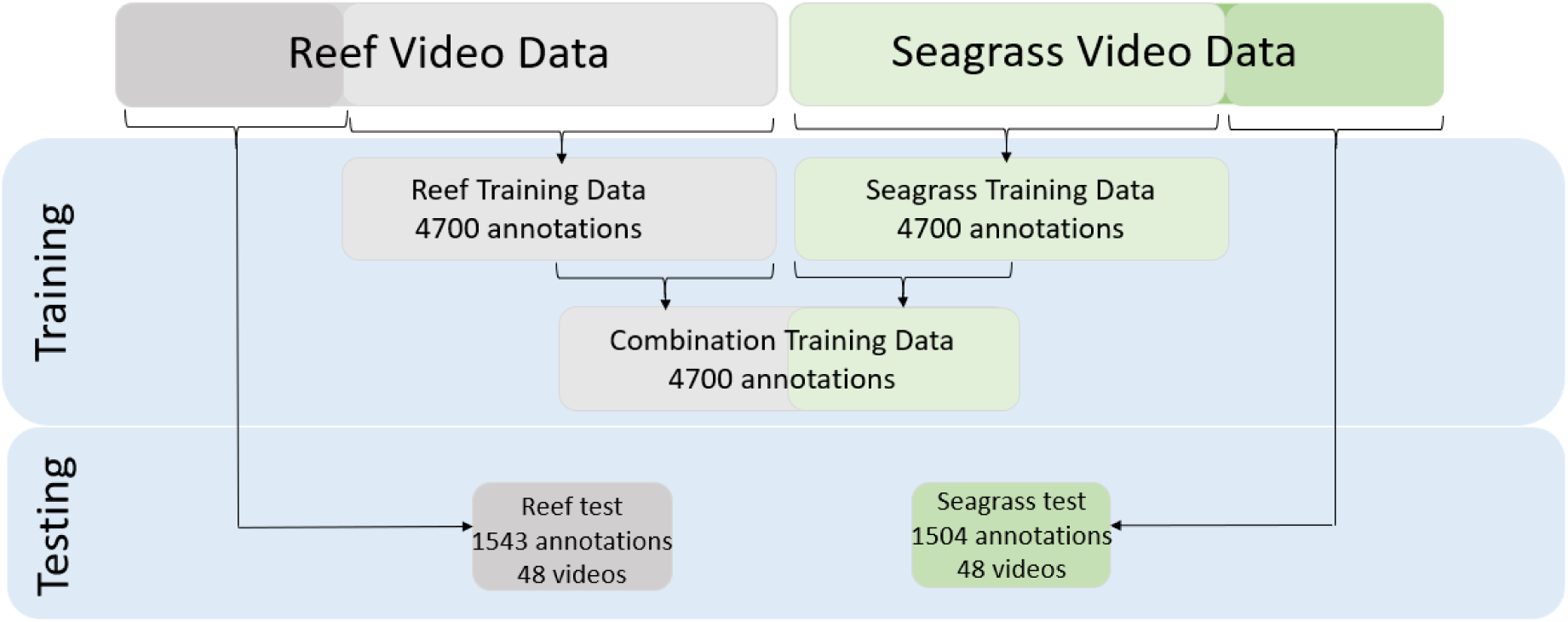
Trimmed videos from the original dataset from two habitats (seagrass and reef) were split into two training sets, with a random selection of half of the annotations from both training sets used to create the combination training set. Two separate tests were created from the remaining datasets from the seagrass and reef habitats.

### Convolutional Neural Network

The object detection framework we used is an implementation of Mask R-CNN developed by Massa & Girshick (2018). Model development was conducted using a ResNet50 configuration, pre-trained on the ImageNet-1k dataset. Model training, testing and prediction tasks were conducted on a Microsoft Azure Data Science Virtual Machine powered by an NVIDIA V100 GPU. Overfitting was minimised by using the early-stopping technique (Prechelt 1998).

### Performance measurements

We tested the model’s performance both for object detection and measuring fish abundance. Object detection performance was determined for each test as the mean average precision 50 value (mAP50, Everingham et al. 2010). This is the ability of the algorithm to accurately fit a segmentation mask to at least 50% of the ROI. Fish abundance performance was tested using MaxN, the maximum number of fish of the target species in any frame in a video, the most widely reported measure in ecological studies (Whitmarsh et al. 2017). Performance on MaxN was calculated by an F1 score; the harmonic mean of precision and recall (Goutte and Gaussier 2005). True positives (the number of correctly identified luderick), false negatives (a luderick was present, but the algorithm did not detect it) and false positives (no luderick present, but the algorithm detected one) are all considered when calculating precision and recall (Buckland and Gey 1994).

## Results

In terms of object detection performance, the seagrass and reefs tests did not perform as well when trained on footage exclusively from the other habitat (Table 1). However, mAP50 performance for tests trained on footage from the same habitat or from a combination of habitats were all high (> 89.9%) and within 4.2% of each other (Table 1), indicating that the algorithm accurately fitted segmentation masks around luderick. Both mAP50 test scores for the combination trained model came within 1% of the same-habitat trained models.

**Table 1.** Summary of model performance. Object detection performance reported as mAP50. Abundance measuring performance reported, overall, as F1 score, denoting how well the model estimated MaxN per video, along with component statistics. Ground truth is the number of fish in videos, and Total fish is the number detected by the model (calculated by the number of true positives and false positives). The F1 denotes how well the model estimated the MaxN per video within each test.

|  |  | Object<br>detection<br>performance | Abundance measurement performance |  |  |  |  |  |
| --- | --- | --- | --- | --- | --- | --- | --- | --- |
| Test | Training | mAP50 | True<br>positive | False<br>positive | False<br>negative | Total<br>fish | Ground<br>truth | F1<br>score |
| Seagrass |  |  |  |  |  |  |  |  |
|  | Seagrass | 91.7 | 99 | 22 | 6 | 121 | 105 | 87.6 |
|  | Reef | 73.3 | 75 | 6 | 30 | 81 | 105 | 80.6 |
|  | Combination | 92.4 | 102 | 18 | 3 | 120 | 105 | 90.7 |
| Reef |  |  |  |  |  |  |  |  |
|  | Seagrass | 58.4 | 11 | 86 | 17 | 97 | 128 | 68.3 |
|  | Reef | 86.9 | 120 | 17 | 8 | 137 | 128 | 90.6 |
|  | Combination | 87.8 | 121 | 32 | 7 | 153 | 128 | 86.1 |

In terms of the model’s ability to determine abundance, the overall pattern was that combination training gave excellent results, as did training singularly on the habitat being tested (Table 1). For the seagrass test, combination training slightly improved on seagrass training (F1 90.7% vs 87.6%, respectively), whereas for the reef test reef training was slightly better than combination training (F1 90.6% vs 86.1%, respectively). The combination training achieved the lowest number of false negatives for both tests, and most closely enumerated the true number of luderick (Table 1). In both tests, the model trained on the opposite habitat gave clearly the worst performance.

## Discussion

Deep learning models trained on a combination of habitats produced the best object detection (mAP50) and either the best or nearly as good abundance estimates (F1). As expected, models also performed very well when tested on the singular habitat they were trained on. They were much poorer at both object detection and abundance estimates when tested on the opposite habitat to that train on. Combination training also had the lowest number of false negatives and most closely enumerated the true number of luderick. The reef dataset, however, had the lowest number of false positives for both tests, suggesting that it was able to account for other environmental factors that could have been confused as luderick, such as other similar looking fish species. In general, the reef footage is comprised of a more complex background, with regularly changing lighting conditions on the substrate as well as greater fish species richness than in seagrass datasets. While the seagrass datasets contain images with comparatively low background complexity, the water was often more turbid and picture clarity was thus affected (Figure 2). These differences between habitats may explain both why models trained on the singular opposite habitat performed poorly, and why the seagrass-trained model performed particularly poorly when tested on reef.

**Fig. 2.**
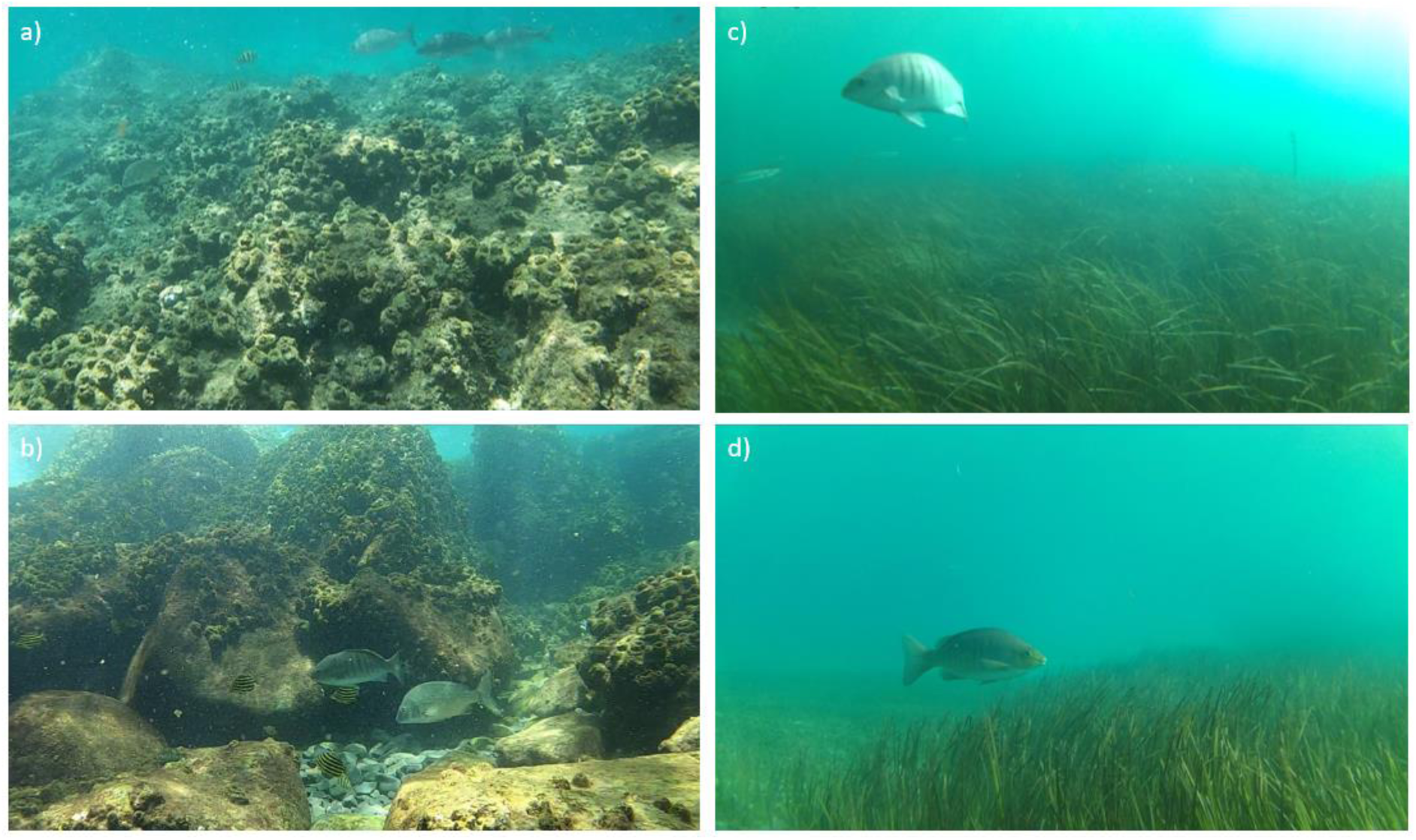
Examples of luderick in test footage from reef (a, b) and seagrass (c, d) habitats, highlighting the diversity of environmental complexities and picture clarity.

To maximise the effectiveness of monitoring and the reliability of analyses, these algorithms must prove robust across habitat types, and often also across significantly separated locations. We have previously shown that deep learning models can have equally high performance in seagrass habitat from a different estuary than where the training footage was taken (Ditria et al. 2019). The transferability of algorithms from one habitat type and location to novel habitat types and locations strengthens this as an alternative option to manual analysis. However, this is not always the case. For example, Xu and Matzner (2018) found that for three different sites, a deep learning model for fish detection trained on two sites, and tested on the third, did not perform as well as those trained and tested on the same sites. This low transferability may have been due to variable water clarity making it difficult to detect fish in video footage (Xu & Matzner 2018). Further testing of fish detection algorithms across habitats and locations is required; clearly training is improved when some variation in habitats is captured, but new training may not be required for every new habitat or location across a species distribution. An additional advantage of deep learning is that unlike traditional machine learning algorithms, deep learning algorithms are not saturated at higher volumes of data, so additional training data will generally improve the overall output performance of the existing dataset (Moniruzzaman et al. 2017; Sarwar et al. 2019). This advantage of deep learning, along with our results, suggests that adding training from newly encountered habitats could be used to continuously improve monitoring results.

Although environmental issues such as different backgrounds, turbidity, lighting and colour hue are not dissimilar to those faced by humans when identifying fish from videos, deep learning algorithms can out-perform humans when faced with ambiguous images (Villon et al. 2018; Ditria et al. 2019). Mask R-CNN can “learn” that the unselected confounding background pixels are not the region of interest, and does not require complex pre-processing of images for background subtraction (Massa and Girshick 2018). Comparing model performance against humans, as well as additional testing in different locations, habitat and time periods would aid in strengthening the idea that these deep learning algorithms are a viable alternative to current manual analysis across large temporal and spatial scales.

Given our rapidly changing world, using robust and flexible deep learning algorithms to monitor and track changes in species occurrence and abundance across entire spatial distributions is important. Efficient monitoring of luderick, for example, could benefit coastal ecology science. Luderick are a significant component of the total fish biomass on temperate reefs on the east coast of Australia, and as algal grazers are considered very important determinants of algal growth rates (Ferguson et al. 2015). Abundance data for luderick are currently very limited and geographically patchy (Abrantes et al. 2015). Still, there is a suggestion that their numbers have declined substantially at the northern (warmer) end of their range in southeast Queensland in recent decades (Pollock 2017). Waters along the south east coast of Australia are experiencing a warming rate over three times the global average (Ridgway 2007), leading to the tropicalisation of historically temperate reefs (Hobday and Pecl 2014; Vergés et al. 2018), and southward range shifts on Australia’s east coast have been fully documented for several other fish species (Townhill et al. 2019). A southward shift in luderick distribution will reduce their role as a key grazer at the current northern limit of their distribution. Monitoring changing range shifts has become a necessary task for management and conservation of functional habitats and implementing deep learning solutions to analyse the abundance of data available is promising. Deep learning algorithms, when trained across a variety of habitat types, could assist in ecological monitoring such as tracking these distribution shifts and changes in population sizes for a range of fish species.

The emergence of deep learning as an accessible and alternative method to manage and extract information from large volumes of raw video footage. The use of a diverse training data set consisting of different habitats and a range of environmental conditions proved to be the most robust and flexible model when analysing footage from different habitats. These models can continually be added to without any adverse effects on performance. Deep can offer rapid data analysis of a range of monitoring activities with high efficiency, and a high level of accuracy and consistency.

## Funding

RC was supported by a Discovery Project from the Australian Research Council (DP180103124). All authors were supported by the Global Wetlands Project.

## Author contributions statements

ED and RC designed the study. ED and SL conducted the fieldwork. ED and EJ developed the deep learning architecture and user interface. RC provided resources. All authors helped interpret results. ED led the writing of the manuscript, with input from all authors

## Conflict of interest

The authors declare that the research was conducted in the absence of any commercial or financial relationships that could be construed as a potential conflict of interest.

## Notes

### Competing Interest Statement

The authors have declared no competing interest.

